# A Sample Covariance-Based Approach For Spatial Binary Data

**DOI:** 10.1101/2020.07.06.189571

**Authors:** Sahar Zarmehri, Ephraim M. Hanks, Lin Lin

## Abstract

The field of landscape genetics enables the study of infectious disease dynamics by connecting the landscape features with evolutionary changes. Quantifying genetic correlation across space is helpful in providing insight into the rate of spread of an infectious disease. We investigate two genetic patterns in spatially referenced single-nucleotide polymorphisms (SNPs): isolation by distance and isolation by resistance. We model the data using a Generalized Linear Mixed effect Model (GLMM) with spatially referenced random effects and provide a novel approach for estimating parameters in spatial GLMMs. In this approach, we use the links between binary probit models and bivariate normal probabilities to directly compute the model-based covariance function for spatial binary data. Parameter estimation is based on minimizing sum of squared distance between the elements of sample covariance and model-based covariance matrices. We analyze data including *Brucella Abortus* SNPs from spatially referenced hosts in the Greater Yellowstone Ecosystem (GYE).

## 1 Introduction

Many disciplines such as ecology, epidemiology and forestry frequently encounter spatial binary data (see e.g. Lannarilli et al. (2019), Hughes et al. (2011), Heagerty and Zeger (1998), Polson et al. (2013)), Rowlingson et al. (2002), Fortin et al. (2013)). However, most spatial data are limited to one observation or replicate at any spatial location. Having many replicates of spatial data is rare, but recent advances have made collecting spatially referenced genetic data possible in a way that results in thousands of replicated binary spatial observations. Our work develops methods to overcome the challenges caused by having many replicates of spatial binary data.

The field of landscape genetics is “an amalgamation of molecular population genetics and landscape ecology (Manel et al., 2003).” This field connects landscape structure with micro-evolutionary processes by first identifying genetic patterns and then connecting the identified patterns to the landscape. One common spatial genetic pattern is called “isolation by distance” and captures the idea that genetic distance or dissimilarities increases with increasing geographic distances (Broquet et al., 2006). Other patterns include “isolation by resistance” (McRae, 2006), in which genetic dissimilarity is a function of landscape features such as rivers and roads located between spatial sampling locations.

The field of landscape genetics enables the study of infectious disease dynamics by connecting the landscape features with evolutionary changes. Infectious disease spread has been studied in humans and domestic animals widely but empirical studies of bacterial disease dynamics in wildlife are often constrained due to practical challenges such as limited sample sizes. *Brucella abortus* is a bacterium which causes infections (brucellosis) in humans, livestock, and wildlife, and leads to abortions in female ungulates. Infections transmit primarily through direct contact with aborted fetuses, birthing fluids, and placentas. The Greater Yellowstone Ecosystem (GYE) is the only remaining reservoir of *B. abortus* in the U.S. where the bacterium infects wild bison, elk, and occasionally domestic bison and cattle. Understanding the disease spread dynamics could help to prevent significant financial losses in livestock industry (Kamath et al., 2016).

Kamath et al. (2016) consider *B. abortus* data including single-nucleotide polymorphisms (SNPs) that occurred at 1463 genome loci in 237 spatially referenced hosts in the GYE. The samples were collected from elk, bison (wild and domestic), and cattle between years 1985 to 2013 (Kamath et al., 2016). Although many studies have been done to connect evolutionary changes to landscape features, it is common to reduce the genetic data to a pairwise genetic dissimilarity matrix, which may ignore much of the rich information in the SNP data (Hanks and Hooten, 2013; Smouse and Peakall, 1999).

The scientific motivation of this paper is to investigate isolation by distance and isolation by resistance patterns in *B. abortus* SNP data. Quantifying genetic correlation across space can provide insight into the rate of spread of *B. abortus* across the GYE. Different models including Generalized Linear Mixed effect Model (GLMM) with spatially structured random effect (Diggle et al., 1998), Generalized Estimating Equations (GEE) (Albert and McShane, 1995), and autologistic models (Besag, 1972) exist for spatial binary data. We model the SNP data using a GLMM with spatially structured random effects.

The most common approach in spatial statistics for inference on parameters in GLMM with spatially structure random effects is Markov Chain Monte Carlo (MCMC). Because of the size of the data and large number of random effects required in a GLMM (one spatial random field for each of the 1463 loci), estimating model parameters using MCMC is computationally impractical. Some packages in R such as glmmTMB (Brooks et al., 2017) might be used for similar data but they only supports a small selection of covariance matrices. Another approach which has common features with our approach is based off of the lorelogram (Heagerty and Zeger, 1998). The lorelogram measures dependence based on marginal pairwise log-odds ratios. According to Heagerty and Zeger (1998), the lorelogram is an alternative method to the variogram (Cressie, 1993) for binary or categorical data. Heagerty and Zeger (1998) mention that for binary data, the variogram or correlogram is constrained by the mean and propose to use lorelogram instead especially when the mean is non-stationary (Heagerty and Zeger, 1998; Lannarilli et al., 2019).

In this paper, we propose a novel approach to compute the covariance of spatial binary data using numerical approximation of multivariate orthant probabilities. To estimate model parameters, we propose a sample covariance-based method by minimizing the squared distance between the model-based covariance and sample covariance. Our approach is similar to variogram or co-variogram approaches. We also used lorelogram-based approach to estimate model parameters by minimizing the sum of squared differences between model lorelogram and empirical lorelogram. We use *B. abortus* landscape genetic data from Kamath et al. (2016). Our paper differs scientifically from Kamath et al. (2016) primarily in our formal investigation of landscape effects such as elevation and the percent of forested areas on gene flow by quantifying them through isolation by resistance, which were not considered by Kamath et al. (2016).

Aside from the scientific contribution of our paper in investigating isolation by distance and isolation by resistance patterns in *B. abortus* SNP data, the novelty of this paper is more comprehended in the proposed methodology. The methodological contributions are as follows:

- We develop a novel approach for analyzing binary GLMMs with spatially referenced random effects to overcome the computational burden. Our approach is based on computing the model-based covariance for spatial binary data using numerical approximations. We propose to minimize the sum of squared differences between elements of the sample genetic covariance and the corresponding model-based covariance for efficient parameter estimation. Sum of squared differences or squared Frobenius norm is a common method to measure the distance between two matrices. Our approach follows the same logic as variogram approaches which are based on minimizing the difference between a variogram estimator and the model-based variogram (Cressie, 1993).
- Different from the classical variogram approaches where the underlying model typically follows a Gaussian distribution, our approach naturally applies to data generated from more general distributions such as binary data.
- We develop a novel method which uses links between binary probit models and bivariate normal probabilities in order to directly compute the covariance function for spatial binary data with spatially-correlated latent random effects. Our method is general enough to allow for any parametric covariance model for the latent spatial random effects, and we illustrate our method using both geostatistical and autoregressive models for spatial random effects.

In Section 2, we introduce spatial GLMMs and our model for the SNP data. In Section 3.1, we present a Bayesian approach and MCMC algorithm for estimating the model parameters. In Section 3.2, we propose our sample covariance-based approach for parameter estimation. In Section 4, we compare the two approaches on simulated data and show that our sample covariance-based approach is much more computationally efficient than MCMC. In Section 5, we apply our sample covariance-based method to the *B. abortus* data and interpret the results. We conclude with discussions in Section 6.

## 2 A Generalized Linear Mixed Effect Model for SNP Data

The *B. abortus* data, which is available as supplementary information in Kamath et al. (2016), include *L* = 1463 genome loci from *N* = 237 spatially referenced hosts at *S* = 74 unique spatial locations in the GYE. At each of the *L* loci in the *B. abortus* genome, two or three alleles are possible. If any variation from the most common allele in a specific locus occurs, we say there is a SNP at that specific locus. Let *𝒮* = *{s*_1_, *s*_2_, …, *s*_*S*_ *}* represent the set of *S* unique spatial locations, *ℒ* = *{l*_1_, *l*_2_, …, *l*_*L*_*}* represent the set of *L* loci, and 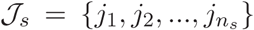 represent the set of *n*_*s*_ individuals in spatial location *s*.

Figure (1) shows the *B. abortus* data related to one of loci *ℒ* and all spatial locations *𝒮* over the map of the GYE. The color of a circle is red if there is a SNP in location *s ∈ 𝒮*, and it is blue otherwise. The size of the circles is proportional to *n*_*s*_– the number of individuals in each spatial location *s ∈ 𝒮*. The map is colored with respect to the elevation, with green indicating high elevation terrain. From this map we can clearly see spatial patterning in the binary genetic data, as locations with SNPs present are clustered together.

**Figure 1:**
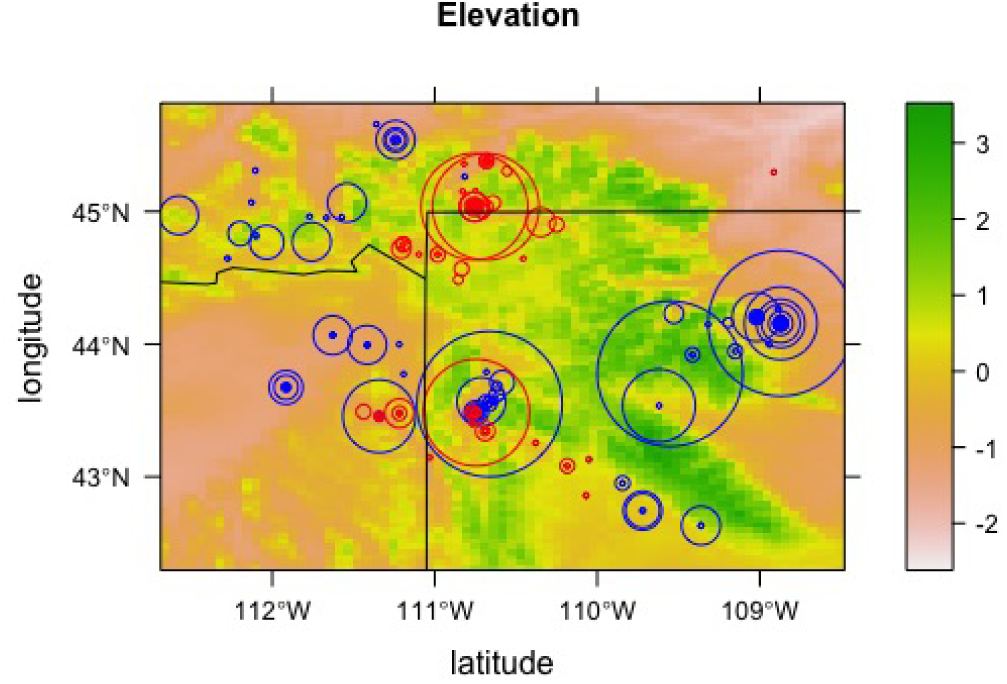
In each location, a circle is red if there is a SNP and it is blue if there is no SNP.

Let ***Y***_*l*_ be a random vector with elements *Y*_*sjl*_ such that *Y*_*sjl*_ = 1 if there is a SNP in locus *l* of genome of individual *j* in spatial location *s* and *Y*_*sjl*_ = 0 otherwise, for *s ∈ 𝒮, j ∈ 𝒥*_*s*_, and *l ∈ ℒ* In this model, we assume that **Y**_*l*_ is independent of **Y**_*l*_ for all loci *l ≠ l′*, and specify a binary probit model for *Y*_*sjl*_. Although in some cases, the independence assumption may not be appropriate for all the available loci, it is appropriate for a set of independent tag SNPs. More details about how to address correlation between loci is provided in Section 5. Following Albert and Chib (1993), the binary probit model can be represented in terms of a latent variable *Z*_*sjl*_:

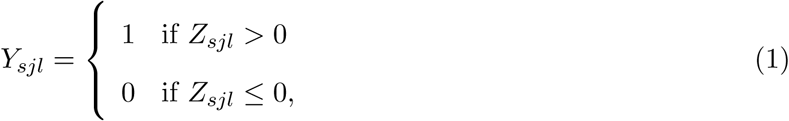

where

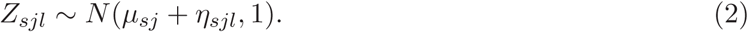

Hence the probability of having a SNP in the genome of individual *j* at spatial location *s* and locus *l* is

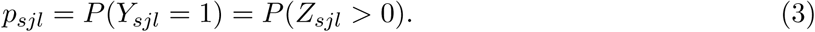

The mean of *Z*_*sjl*_ in (2) consists of two parts: the intercept *µ*_*sj*_ and a spatially structured random effect *η*_*sjl*_ specific to the locus *l*. Depending on the nature of the problem, one can define a varying intercept *µ*_*sj*_ to include landscape or individual features in the mean structure of *Y*_*sjl*_. For example, it is possible to define *µ*_*sj*_ = ***β′x***_***s***_ + *ϵ*_*sj*_ where ***x***_***s***_ is a vector of landscape covariates at location *s* and ***β*** are their associated parameters to estimate. We do not include landscape features in our model for *E*(*Y*_*sjl*_); Doing so would be reasonable if we assumed genetic variation was caused by adaptation rather than drift. However, since our data consist of *B. abortus* isolates obtained from different species (elk, bison, and cattle), we model a different intercept for each species. Thus *µ*_*sj*_ = *µ*_*e*_ if the *B. abortus* was obtained from elk at site *s, µ*_*sj*_ = *µ*_*b*_ if it was obtained from bison, and *µ*_*sj*_ = *µ*_*c*_ if it was obtained from cattle. Larger values of *µ*_*sj*_ and *η*_*sjl*_ make it more likely to observe SNPs in the data. Let ***η***_*l*_ be a random effect vector with elements *η*_*sjl*_ for *s ∈ 𝒮, j ∈ 𝒥*_*s*_, and *l ∈ ℒ* and we assume that ***η***_*l*_ independently follows a multivariate normal distribution:

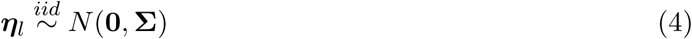

where Σ is a spatially structured covariance matrix.

In this paper, we consider two different models for the covariance matrix Σ. In the first model, which is based on the “isolation by distance” approach (Broquet et al., 2006), Σ is parameterized using an exponential covariance function with

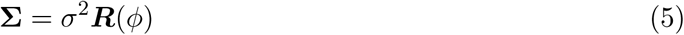

where σ^2^ is the partial sill and ***R***(*ϕ*) is a *N × N* positive semidefinite correlation matrix. For all *s, s′ ∈ 𝒮, j′ ∈ 𝒥*_*s*_, and *j ∈ 𝒥*_*s′*_ let

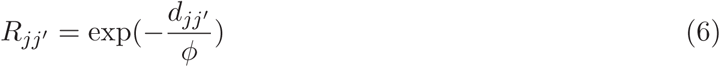

where *d*_*jj′*_ is the Euclidean distance between individual *j* in location *s* and individual *j′* in location *s′* and *ϕ >* 0 is a range parameter.

Our goal is to estimate parameters *µ*_*sj*_, σ^2^, and *ϕ*. The spatial range parameter *ϕ* is of particular interest as it controls the correlation between genetic material across space.

In the second model, we consider a covariance matrix that captures landscape effects on gene flow. The migration of genetic variation from one population to another is called gene flow. We refer to this model as the landscape covariance model. This model is related to the “isolation by resistance” approach (McRae, 2006). McRae (2006) proposed a new method to predict the effect of landscape structure on genetic differentiation called “isolation by resistance”, in which how gene flow behaves in the landscape is very much similar to the effective conductance (reciprocal of resistance) between nodes in an electric circuit. In this framework, one imagines a spatial domain as a graph of spatial nodes where the edge weights are proportional to the rates of random walk between nodes. The rates of random walk in this graph are analogous to effective conductance between nodes in a circuit. Similar to an electric circuit where the effective conductance between two nodes increases by making additional connections, level of gene flow increases if additional parallel movements of genes are allowed. Unlike “isolation by distance” methods which account for landscape effects that increase dispersal or act as barriers by adjusting the Euclidean distance, the “isolation by resistance” model is more flexible to integrate subtle effects of spatial heterogeneity into genetic variation (McRae, 2006).

Hanks and Hooten (2013) showed that isolation by resistance can be seen as a particular form of spatial autoregressive model. Hanks (2017) expands on this to allow for directional bias in random walk rates, or equivalently in gene flow rates. Following Hanks (2017), we specify a covariance matrix which corresponds to the stationary distribution of a spatio-temporal random walk process for gene flow. The spatial covariance matrix Σ is of the form

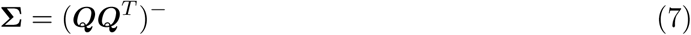

where (***QQ***^*T*^)^−^ is the generalized inverse of ***QQ***^*T*^ and ***Q*** is the infinitesimal generator of the random walk process:

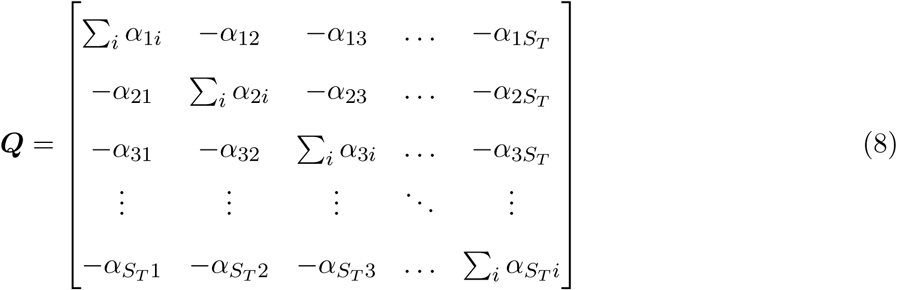

where *S*_*T*_ is the total number of spatial locations (grid cells in a regular lattice) and *α*_*ss′*_ is the random walk rate between locations *s* and *s′*.

We model the random walk rate *α*_*ss*_ in (8) as:

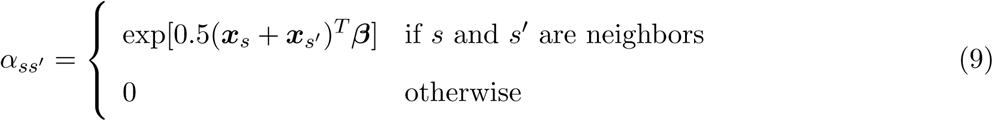

where ***x***_*s*_ and ***xs*** represent the landscape properties of neighboring locations *s* and *s′*, for example elevation at each location, and ***β*** is a vector of random walk parameters.

This model allows for inference on how intervening landscape features (e.g. mountain ranges or rivers) affect spatial correlation. Positive values of *β*_*k*_ imply high gene flow in terrain with large values of *x*_*k*_ while negative values imply low gene flow.

The *B. abortus* data includes observations of *N* = 237 individuals at *S* = 74 unique spatial locations. We have landscape features at *S*_*T*_ = 6525 grid cells with *B. abortus* samples coming from *S* of these cells. Hence out of *S*_*T*_ locations, we have multiple observations in a small subset of them. The precision matrix ***QQ***^*T*^ is typically sparse but its generalized inverse is usually dense. To avoid the computation of (***QQ***^*T*^)^−^ when *S*_*T*_ is large, we use an approach similar to that of Hanks and Hooten (2013). Based on this approach, we expand (7) as

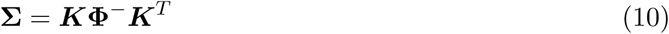

where **Φ** is the precision matrix of uniquely observed locations and ***K*** is a *N × S* matrix with *K*_*js*_ = 1 if *j ∈ 𝒥*_*s*_ for *s ∈ 𝒮* and *K*_*js*_ = 0 otherwise. **Φ** can be obtained efficiently by partitioning ***QQ***^*T*^ and calculating the Schur complement (Harville, 2008). For more details, see Hanks and Hooten (2013).

## 3 GLMM Parameter Estimation

Estimating parameters in these two spatial models will provide insight into the spatial scales at which we expect to find correlated *B. abortus* SNPs. The most common approach in spatial statistics for inference is Bayesian inference (Osborne et al., 2001; Hooten et al., 2003; Gelfand et al., 2005; Christensen et al., 2006; Latimer et al., 2006). However, given the size of our data (*L* = 1463 loci), using a Bayesian approach and fitting the model by MCMC may be computationally impractical. In section 3.2, we will propose a novel approach which is the main contribution of our paper for inference on parameters in the above models when we have hundreds of locations and thousands of loci. Our approach is similar to variogram approaches but, unlike classical variogram approaches in which the underlying model follows Gaussian distribution, we model binary data by incorporating binary GLMMs with spatially referenced random effects and directly compute the covariance function for spatial binary data using numerical approximation.

### 3.1 Bayesian Approach

We first consider a Bayesian approach to estimate the parameters *µ*_*sj*_, σ^2^, and *ϕ* in the exponential covariance model. The joint posterior distribution of the parameters is

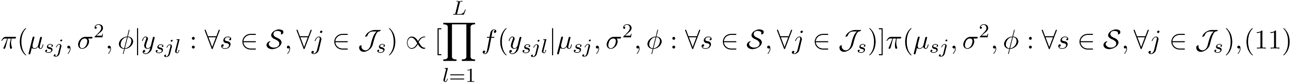

where *f* (*y*_*sjl*_|*µ*_*sj*_, σ^2^, *ϕ*: ∀*s* ∈ S, ∀*j ∈ J*_*s*_) represents the likelihood function of the data related to locus *l* and *π*(*µ*_*sj*_, σ^2^, *ϕ*: ∀*s* ∈ S, ∀*j ∈ J*_*s*_) is the joint prior distribution of the parameters.

At each iteration of the MCMC algorithm, we need to sample *Z*_*sjl*_ and ***η***_*l*_ for all *s ∈ 𝒮, j ∈ 𝒥*_*s*_, and *l ∈ ℒ*, as well as the parameters *µ*_*sj*_, σ^2^, and *ϕ*. Based on (1) and (2), full-conditional samples of *Z*_*sjl*_ are draws from a truncated normal distribution where the upper and lower limits are identified by the value of *Y*_*sjl*_. For the other variables, we need to specify the prior distributions. Following Schliep and Hoeting (2015), we specify conjugate priors for ***η***_*l*_, *µ*_*sj*_, and σ^2^ as 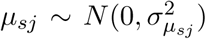, ***η***_*l*_ ∼ *N* (**0**, *Σ*) respectively for each *l ∈ {*1, …, *L}*, where Σ follows the structure in (5) and (6), and σ^2^ ∼ *IG*(*α, β*), and use Gibbs updates to sample from the full conditional distributions. For *ϕ*, we specify a uniform prior as 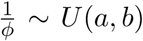 where *a >* 0 and use Metropolis-Hastings (MH) updates. Algorithms (1) and (2) in the appendix A provide additional detail on the MCMC algorithm for the exponential model.

The Bayesian approach for the landscape covariance model is similar to the one for the exponential model. The joint posterior distribution of the parameters is

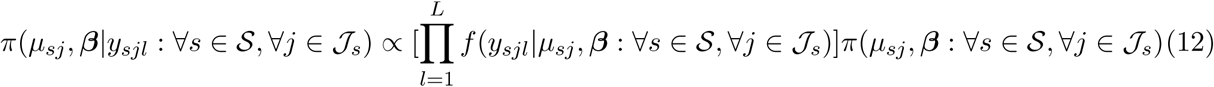

The MCMC sampler is also similar to the sampler for the exponential model described above. The only differences are in the full conditional distributions of ***η***_*l*_ and the covariance parameters. Here, the conjugate prior for ***η***_*l*_ is ***η***_*l*_ ∼ *N* (**0**, *Σ*) where Σ is now coming from (7). The covariance parameters we need to estimate are ***β*** instead of σ^2^ and *ϕ*. We specify a prior 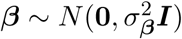 and use Metropolis-Hastings updates.

### 3.2 Sample Covariance-based Approach

The MCMC algorithm above is computationally taxing, as we have *L* = 1463 latent random effects ***η***_***l***_. We propose a sample covariance-based approach to compute the model based covariance by numerical approximation and estimate model parameters by comparing the model based covariance of ***Y***_*l*_ with the sample covariance.

Our approach for estimation of model parameters is related to the least squares fitting of variogram models (Cressie, 1993). Using least squares methods is common in many disciplines and can be interpreted as minimizing the distance between two matrices. In terms of matrix calculation, it is equivalent to minimizing the squared Frobenius norm of the difference between two matrices. As an example in finance, Higham (2002) used matrix norms to find the nearest correlation matrix of stocks. In our approach, we estimate the parameters of model-based covariance by minimizing the sum of squared differences between the model-based and sample covariance matrices where both are proper covariance matrices.

The main methodological contribution of this paper is in the proposed approach which uses the links between binary probit models and bivariate normal probabilities to directly compute the covariance function for spatial binary data with spatially correlated latent random effects. In classical variogram approaches, the underlying model typically follows Gaussian distribution, whereas in this paper, we develop methods for spatial probit models as we are dealing with binary data. The lorelogram is another similar approach to estimating parameters from binary data. To estimate model parameters using the lorelogram, Heagerty and Zeger (1998) proposed using a pair of estimating equations for parametric models. In this paper, we use the least squares fitting of sample and model-based lorelograms to make a comparison with our proposed approach.

Since SNP data are often of very high dimension (we have 1463 loci in our *B. abortus* genetic data), the sample covariance of the data provides an accurate estimator of the true covariance function when the loci are independent. However, SNPs found by whole genome sequencing are often dependent. If the independence assumption is violated, it is possible to find a subset of approximately independent SNPs and use the proposed sample covariance-based approach. More details of this approach are available in section 4. As we model SNP data using a GLMM, it is difficult in most cases to write out the covariance function as the spatial random effects ***η***_*l*_ are applied to the linear predictor in the GLM. However, we present a fast and highly accurate numerical approximation to the covariance function for a binary GLMM under the probit model (1)-(2). This approximation is valid for any covariance function of the spatial random effects ***η***_*l*_, and makes computing the covariogram of binary data fast and reliable.

Using (2) and (4), we can write the probability of having a SNP at genome locus *l* of an individual *j* at spatial location *s* as

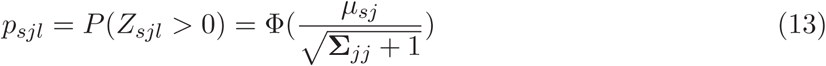

for all spatial locations *s ∈ 𝒮*, individuals *j ∈ 𝒥*_*s*_, and loci *l ∈ ℒ*, where Σ_*jj*_ represent the diagonal elements of the *N × N* spatial covariance matrix Σ of ***η***_*l*_.

Based on our model assumptions, for any *l ∈ ℒ*, the elements of **Y**_*l*_ are correlated with each other because of their spatial dependencies, but they are independent from the elements of **Y**_*l′*_ for any other locus *l′* ≠ *l*. Let Σ_*Y*_ represent the covariance matrix of ***Y***_*l*_ and (**Σ**_***Y***_)_*jj′*_ represent the covariance between *Y*_*sjl*_ and *Y*_*s′j′l*_ which are the elements of ***Y***_*l*_ associated with individuals *j* and *j*′ in spatial locations *s* and *s′* respectively. Based on (2) and (13), we can obtain (**Σ**_***Y***_)_*jj′*_ by

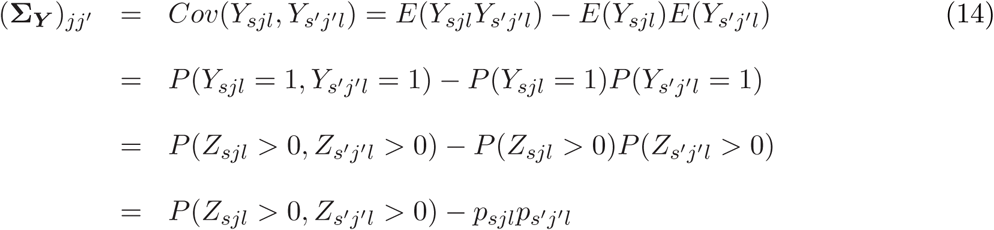

where *P* (*Z*_*sjl*_ *>* 0, *Z*_*s′* *j′l*_ *>* 0) represents the bivariate normal probability that *Z*_*sjl*_, *Z*_*s′* *j′l*_ lies in the 1st quadrant.

Considerable research has been devoted to multivariate normal distribution computations (Genz and Bretz, 2009). We use the numerical method proposed by Drezner and Wesolowsky (1990) and approximate the bivariate normal probability in (14) by the integral

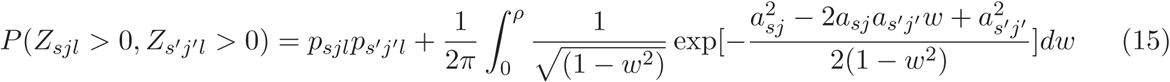

where 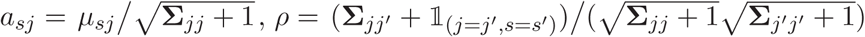, and 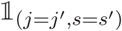 is an indicator function such that 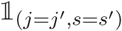 when *sj* = *s′j′* and 0 otherwise.

Substituting (15) in (14), we get

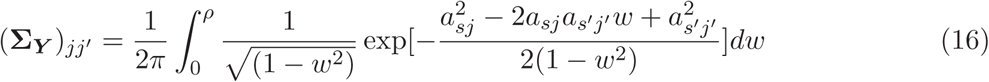

for any *l ∈ ℒ*. Thus, the covariance between any pair of spatially referenced binary observations can be written as a 1-dimensional integral, with the upper limit of the integral (*ρ*) being the correlation of the latent variables *{Z*_*sjl*_, *Z*_*s′j′l*_*}*. This makes directly computing Σ_***Y***_ computationally efficient.

To estimate model parameters, we will compare Σ_***Y***_ with the sample covariance. Let 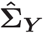 represent the sample covariance of ***Y***

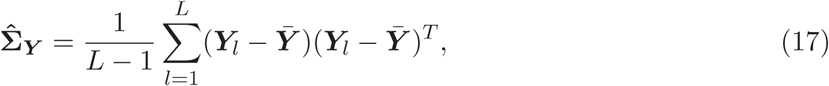

where 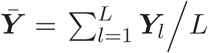.

If the independence assumption is valid, then the consistency of the sample covariance matrix 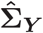 is easy to prove. However, if the assumption is violated, the sample covariance is a biased estimator of the true covariance. The proof is included in appendix B.

For the exponential covariance model, we propose to estimate *µ*_*sj*_, σ^2^, and *ϕ* by minimizing the sum of squared differences between the elements of Σ_***Y***_ and 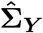 as follows

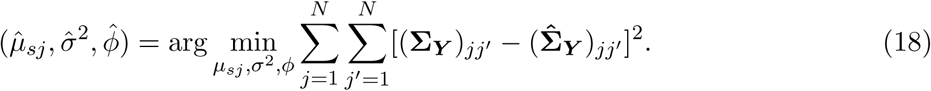

Using the same approach for the landscape covariance model, we estimate model parameters as

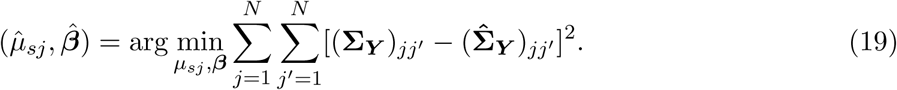

We use the numerical optimization function “optim” in the R statistical computing environment (R Core Team, 2019) to estimate the model parameters.

This sample covariance-based approach for binary spatial models is the main contribution of this work. It is similar to variogram-based approaches, but we have extended these approaches to binary spatial data through providing a form of the model covariance in (16) which is flexible enough to allow for any covariance in the linear predictor of the spatial GLMM and allows for computationally efficient evaluation of the model covariance.

### 3.3 Lorelogram-based Approach

Heagerty and Zeger (1998) proposed the lorelogram as an alternative to variogram approaches for binary or categorical data. The lorelogram, which is the marginal pairwise log-odds ration, is defined as

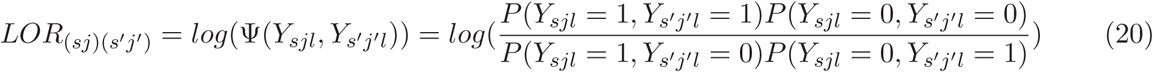

where *LOR*_(*sj*)(*s′j′*)_ represents the lorelogram between *Y*_*sjl*_ and *Y*_*s′j′l*_.

Using equation (15), we can write *LOR*_(*sj*)(*s′j′*)_ as

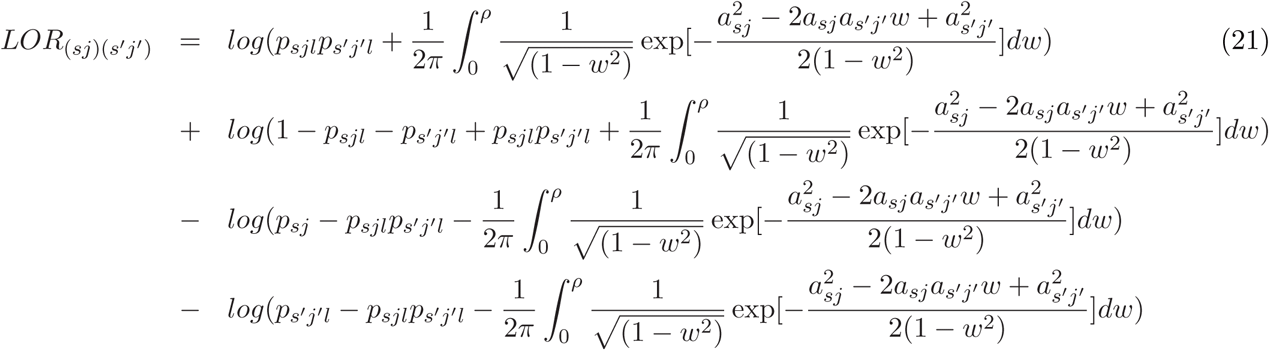

Note that *LOR*_(*sj*)(*s′j′*)_ might be undefined or goes to negative infinity if the probability inside the *log* function goes to zero. This causes some issues in numerical computations and one needs to add some error term to avoid these issues. We added an error term *e*_*lol*_ = 0.1 to all the probabilities inside the *log* function.

For any *s, s′ ∈ 𝒮* and *j, j′ ∈ 𝒥*, let *N*_*cc′*_ represents the number of times the pair (*Y*_*sjl*_ = *c, Y*_*s′j′l*_ = *c′*) appears in the data over all *l ∈ ℒ* where *c, c′ ∈ {*0, 1*}*. Then the empirical lorelogram (Lannarilli et al., 2019) is

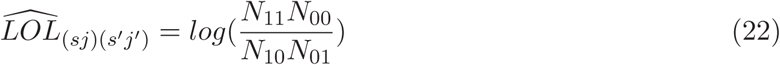

We note that empirical lorelogram is undefined or negative infinity when the denominator or numerator in (22) is 0. To fix this issue, we need to add small error term *e*_*lol*_ to all *N*_*cc′*_ for *c, c′ ∈ {*0, 1*}*. In our estimation using the lorelogram-based approach, we set 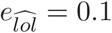.

To estimate the model parameters for the exponential covariance model and landscape covariance model respectively, we minimize the sum of squared differences between model lorelogram and empirical lorelogram as follows

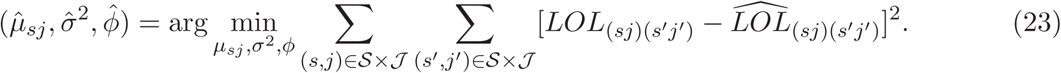

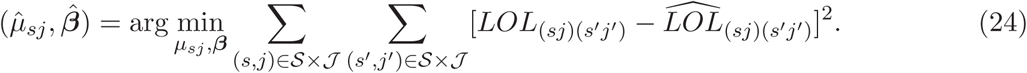

## 4 Simulation Study

In this section, we provide a simulation study comparing the sample covariance-based approach to the Bayesian approach and lorelogram approach on data simulated from the exponential covariance model. Since SNPs found by the whole genome sequencing are dependent, we generate sets ot dependent data so that our simulation study resembles genetic data. Since our estimation approach is based on independent assumption of SNPs, we propose a method to find a subset of approximate independent SNPs first and then apply different estimation approaches.

For any two loci, assume the possible alleles at the first locus are *A* and *a*, and the possible alleles at the second locus are *B* and *b*. A non-random association of alleles at two or more loci is called linkage disequilibrium (LD) (Lewontin and Kojima, 1960) and it depends on the quantity *D*_*AB*_ = *p*_*AB*_ − *p*_*A*_*p*_*B*_ where *p*_*AB*_ is the frequency of gametes at two loci carrying the pair of alleles *A* and *B*, and *p*_*A*_ and *p*_*B*_ are the frequencies of those alleles (Slatkin, 2008). For diallelic loci (which is the case in our data), *𝒟* = *p*_*AB*_*p*_*ab*_ − *p*_*Ab*_*p*_*aB*_. Although *𝒟* completely characterizes the extent of non-random association of alleles *A* and *B*, it is not the best statistic when comparing LD for different pairs of loci (Slatkin, 2008).

To predict one locus with the other one with high accuracy, we need to find the squared correlation coefficient between the two loci, *r*^2^. If *r*^2^ = 1 and we know the allele at locus one, we can predict the allele at locus 2 (Laird and Lange, 2010). *r*^2^ is a commonly used statistics to quantify LD (Slatkin, 2008) and can be calculated as follows:

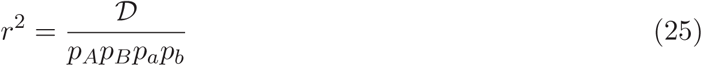

In this paper, we used *r*^2^ to quantify LD as our purpose is to identify a set of independent tag SNPs which can then be used to consistently estimate the spatial covariance matrix. We used *r*^2^ as a measure of similarity to cluster loci. We used hierarchical clustering as implemented in the “hclust” function available in R (R Core Team, 2019) to cluster loci based on their dissimilarity (1 − *r*^2^) and according to the clustering method “average” which stands for unweighted average linkage clustering. Loci with the distance less than 0.8 (which implies *r*^2^ *>* 0.2) were put together in the same cluster. In each cluster, we randomly picked one of the loci as the representative of that cluster and used it in our SNP data analysis. In this way, we can make sure the SNPs used in our analysis are approximately independent.

To generate dependent SNP data, we used a multivariate exponential model. This model is based on the so-called multivariate Matern model (Gneiting et al., 2010). To generate dependent SNPs data, assume for the genome sequence of each individual *j ∈ 𝒥* at locatios *s ∈ 𝒮* we have *{c*_1_, …, *c*_*K*_ *}* clusters of SNPs with size *{m*_1_, …, *m*_*K*_ *}*. In each cluster, the SNPs are dependent while two SNPs from two different clusters are approximately independent. There is also spatial dependencies across locations *s ∈ 𝒮* at each locus *l ∈ ℒ* where the spatial dependencies come from an exponential model. For each cluster of SNPs, our goal is to construct multivariate exponential covariance matrices of sizes *S × m*_*k*_ for *k ∈ {*1, …, *K}*. We can write the multivariate exponential covariance matrix with cross-correlation coefficient *ρ* as follows:

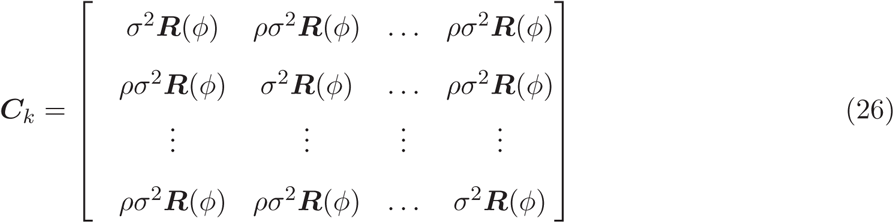

where ***R***(*ϕ*) is given in (6).

For simulating data, we assume a common intercept *µ* for all observations and each spatial location *s ∈ 𝒮* is only associated to one individual. For each cluster *c*_*k*_ with *k ∈ {*1, …, *K}* and all spatial locations *s ∈ S*, let ***Y***_*k*_ be a vector of size *S × m*_*k*_ with elements *Y*_*sk*_ from the following model:

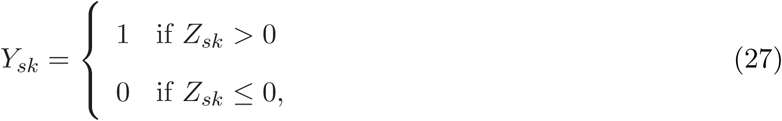

where

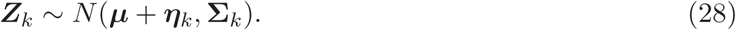

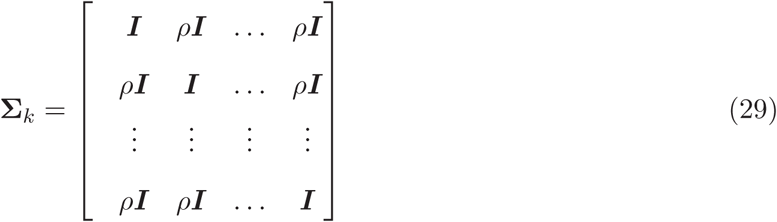

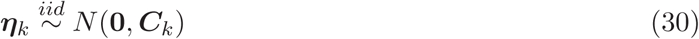

where ***Z***_*k*_ and ***η***_*k*_ are vectors of size *S × m*_*k*_ for all *k ∈ {*1, …, *K}*, ***C***_*k*_ is multivariate exponential covariance matrix defined in (26), and ***I*** is the identity matrix of size *S × S*. Equations (27)-(30) provide a framework to simulate spatial binary data with correlated SNPs. If we pull out one SNPs from each cluster of SNPs, then the resulting set of tag SNPs are independent and follows our model (1)-(4).

Model (27)-(30) allows us to generate clusters of dependent SNPs. According to this model, we have dependency between SNPs and across space in each cluster but different clusters are independent from each other. Our goal is to first cluster the simulated data using the provided tag SNP method and pick a tag SNP as a representative of each cluster to construct an approximately independent set of tag SNPs. We then use this set of approximately independent tag SNPs to estimate model parameters *µ*, σ^2^, and *ϕ* using (18).

Since MCMC is computationally taxing, we generated smaller data sets to compare MCMC with the sample-covariance based approach and lorelogram approach. For our simulation study, we first generated 200 random spatial locations in the unit square. We generated *K* = 20 and *K* = 100 clusters of approximately independent SNPs where each cluster is of size *m*_*k*_ = 5. The cross-correlation coefficient is picked randomly from a *Uniform*(0.8, 1) random variable and is fixed among all the simulations in this section. The randomly selected value is *ρ* = 0.937. We also fixed the values of *µ* = −1, σ^2^ = 2, and *ϕ* = 0.1 to generate the data according to model (27)-(30). The computations are performed on a heterogeneous cluster that consists of multiple node-types connected to a common file system. The node configurations are 2.2 GHz Intel Xeon Processor, 24 CPU/server, 128 GB RAM, and 40 Gbps Ethernet. We parallelized the sample-covariance based approach and lorelogram on 19 processors with 16 GB of RAM.

Table (1) compares the results of 3 methods for *K* = 20 and *K* = 100 with *m*_*k*_ = 5. Therefore the total number of loci in the simulated data is *L* = 100 and *L* = 500 respectively. We then applied our method of choosing tag SNPs to each of these data sets. Based on our results, the tag SNPs method selected 20 tag SNPs out of *L* = 100 and 100 tag SNPs out of *L* = 500. For the sample-based covariance and lorelogram-based covariance approaches, we calculated confidence intervals (CIs) using an empirical bootstrap approach. At each bootstrap iteration, we randomly sampled with replacement from the set of tag SNPs chosen by our tag SNPs method. The size of each bootstrap sample is equal to the size of tag SNPs data set. Note that our bootstrap method assumes independent SNPs and by conducting the method on the tag SNPs data set, ensures that the data are approximately independent. In Table (1), *COV* stands for the sample-covariance based method and *LOL* stands for the lorelogram-based method. For more details of the MCMC algorithm, see algorithms (1) and (2) in the appendix A. For MCMC, we set 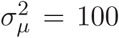, *α* = 3, *β* = 8, *a* = 1, and *b* = 1000 and ran the algorithm for 30000 iterations and discarded the first 10000 as burn-in.

**Table 1:**
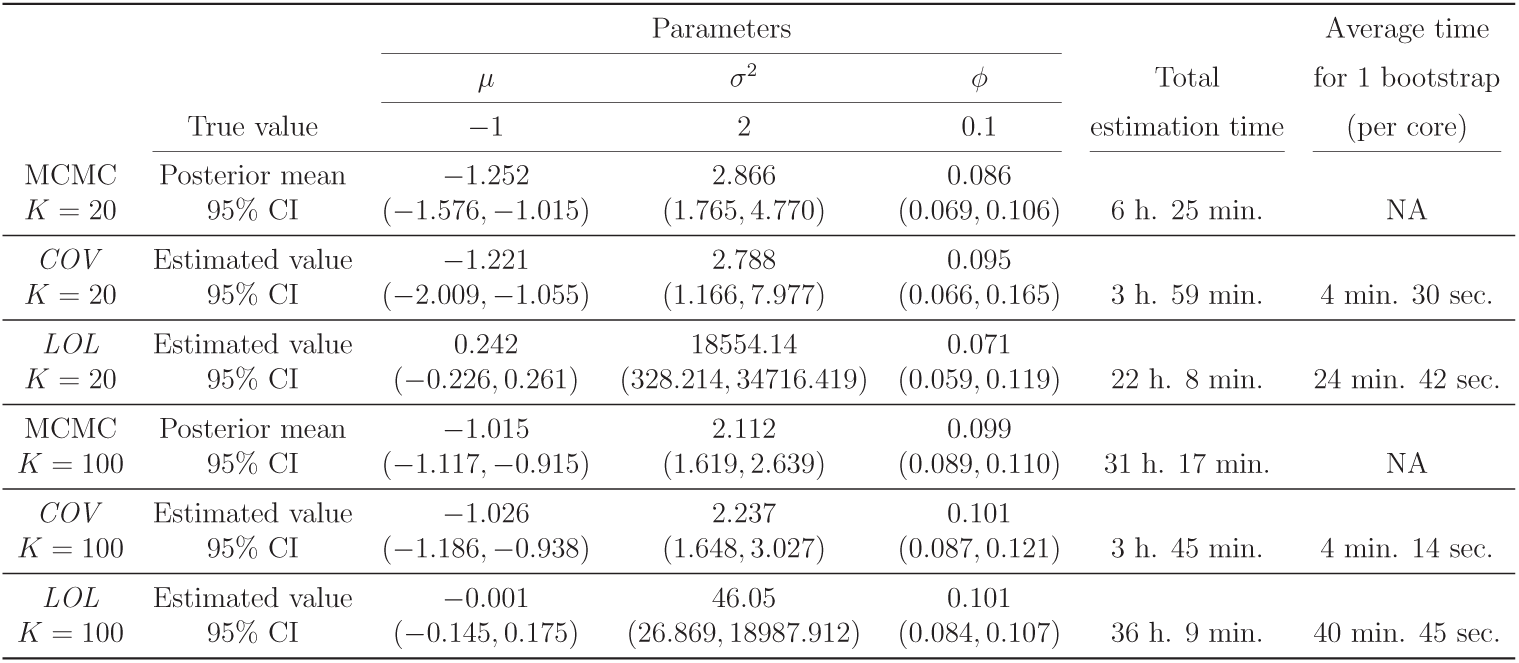
Simulation study results for *K* = 20, and *K* = 100 with *m*_*k*_ = 5. 95% CI represents 95% credible intervals for MCMC and 95% confidence intervals for COV and LOL.

It is evident from Table (1) that both MCMC and *COV* are more successful in estimating the parameters compared to *LOL*. When the number of independent SNPs are small, both MCMC and *COV* methods tend to underestimate *µ* and the provided CIs for σ^2^ cover a wider range of values. As the number of independent SNPs increases, the CIs become tighter and all of them include the true parameter values. This is expected since by increasing the number of SNPs, more information is available for the analysis. Among the two methods, MCMC seems to provide tighter CIs. However, the time needed to estimate the parameters increases linearly as we increase the number of independent SNPs. For example, as we go from *K* = 20 to *K* = 100, the estimation time multiplied by almost 5. So if we want to increase the number of independent SNPs to 1000, the estimation time would be around 314 hours. The time needed for the *COV* method is not affected by the number of SNPs and stays relatively constant. The total estimation time for *K* = 100 is reduced by 1/8 compared to the MCMC. Moreover, one is able to parallel it over more cores to get the results faster. The *LOL* performs very poorly in estimating the two parameters *µ* and σ^2^ but it is able to estimate *ϕ* correctly specially when the size of data increases. The estimation time in the *LOL* is affected by the increasing number of required iterations in the numerical estimation and optimization parts as well as the increase in the number of independent SNPs. However, as we will see in Table (2), as the data size increases and the results become more accurate, the time variation is mainly due to the increasing number of SNPs. We have tried different methods of numerical integration for the LOL, including Gauss-Kronrod quadrature and Simpson method. However, the LOL method is numerically unstable for binary data, and this instability either persisted or leaded to a long convergence time across different numerical approaches for approximating the integrals in (21), and for different values of *e*_*lol*_ and 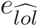. As such, we find that covariogram based approaches are more stable for binary data than are lorelogram based approaches in their current form.

**Table 2:** Simulation study results for *K* = 250, and *K* = 500 with *m*_*k*_ = 5. 95% CI represents 95% confidence intervals for COV and LOL.

|  |  | Parameters |  |  | Total<br>estimation time | Average time<br>for 1 bootstrap<br>(per core) |
| --- | --- | --- | --- | --- | --- | --- |
| | | $\mu$ | $\sigma^2$ | $\phi$ | | |
|  | True value | -1 | 2 | 0.1 |  |  |
| *COV* | Estimated value | -1.042 | 2.08 | 0.102 |  |  |
| *K* = 250 | 95% CI | (-1.136, -0.980) | (1.722, 2.520) | (0.092, 0.114) | 3 h. 51 min. | 4 min. 20 sec. |
| *LOL* | Estimated value | -0.000 | 2.679 | 0.100 |  |  |
| *K* = 250 | 95% CI | (-1.124, 0.001) | (2.305, 3.665) | (0.087, 0.106) | 18 h. 58 min. | 21 min. 9 sec. |
| *COV* | Estimated value | -1.012 | 1.961 | 0.100 |  |  |
| *K* = 500 | 95% CI | (-1.066, -0.969) | (1.734, 2.216) | (0.093, 0.107) | 3 h. 42 min. | 4 min. 10 sec. |
| *LOL* | Estimated value | -1.403 | 1.933 | 0.093 |  |  |
| *K* = 500 | 95% CI | (-1.651, 0.000) | (1.737, 2.714) | (0.086, 0.102) | 28 h. 22 min. | 31 min. 47 sec. |

To check the performance of *COV* and *LOL* method for larger data sets, we simulated two more data sets where *K* = 250 and *K* = 500 with *m*_*k*_ = 5. Therefore, the total number of loci is *L* = 1250 and *L* = 2500 respectively. All other parameters remained the same as before and we used empirical bootstrap method to estimate the CIs. We didn’t perform MCMC for these two sets as it would take around 78 hours for *K* = 250 and 157 hours for *K* = 500. Table (2) includes the results:

From Table (2), the *COV* estimated the parameters more accurately and the CIs include the true parameters values. The time of estimation does not change by increasing the number of SNPs and stays relatively the same. The *LOL* method provided much better results as the size of data increases. The estimation time changes by increasing the number of SNPs but the change is not linear. However, the estimation time is 4 times higher compared to the *COV* method for *K* = 250 and it grows larger as we increase the number of independent SNPs.

Comparing the three methods, the sample-covariance based method performs as good as MCMC for larger data sets while outperforming MCMC in computational speed. To ascertain the CI properties, we found the coverage of *COV* for the case of *K* = 500 with *m*_*k*_ = 5 by generating 100 random data sets using the provided parameter values and constructing the CIs using the bootstrap method. Out of 100 calculated CIs for *µ*, 5 of them did not include the true parameter value *µ* = −1. The same thing is true for the other two parameters σ^2^ and *ϕ*, i.e. 5 out of 100 Cis did not include the true values *sigma*^2^ = 2 and *ϕ* = 0.1.

## 5 *B. abortus* Data Analysis

As we have mentioned before, since the SNPs independence assumption may not be valid for all available loci in the data, we compared two different SNP sets: a set by assuming all available SNPs are independent (call this *set*_*full*_) and a set by first clustering SNPs using linkage disequilibrium (LD) and clustering method “average” in clustering function “hclust” in R statistical computing environment (R Core Team, 2019) and then choosing approximately independent tag SNPs as representatives of clusters (call this *set*_*tag*_).

We now analyze the *B. abortus* data using both covariance models with both sample-based covariance method and lorelogram-based method and compare their results by 10-fold cross validation. As mentioned in Section 1, the data includes SNPs in *L* = 1463 loci from *N* = 237 individuals. The number of unique spatial locations where data were collected is *S* = 74. To find the Euclidean distance between the locations, we used the “fields” package in R (Nychka et al., 2017) and computed the pairwise distance between spatial locations in *𝒮* based on latitude and longitude. For the landscape covariance model, we need to link the observed spatial locations to the landscape raster cells. Using the “raster” package in R (Hijmans et al., 2016), we found the covariate rasters and linked the observed locations to the raster cells. The total number of raster cells is *S*_*T*_ = 6525. The covariates we have considered in the landscape model are an intercept (int), which is 1 for all raster cells, elevation (elev) which has been standardized to have mean 0 and standard deviation 1, and the percent of raster cell that is forested (frst). Figure (2) shows these two covariates over the map of the GYE.

**Figure 2:**
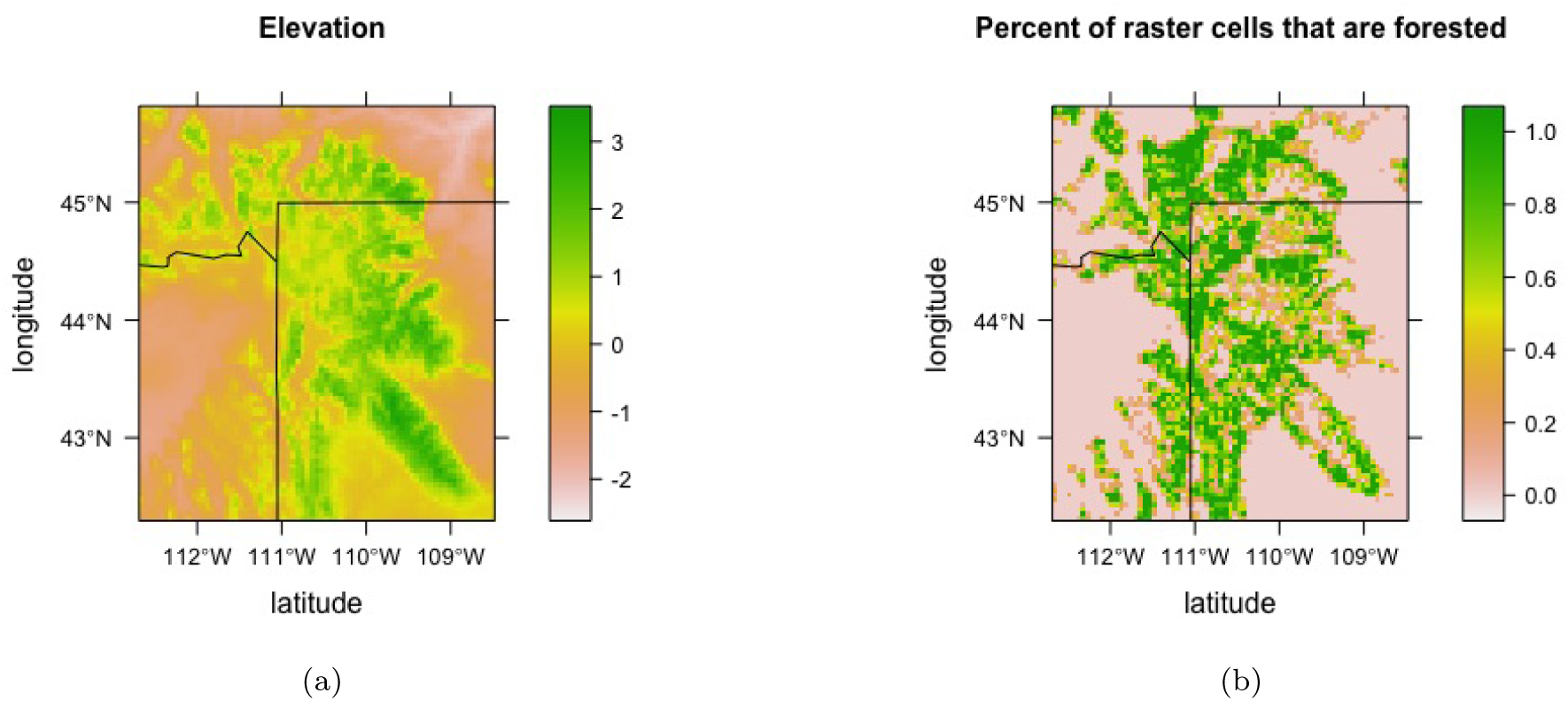
(a) The more green the area, the higher the elevation. (b) The more green the area, the higher the percent of forested cells.

For each covariance function, we modeled the *B. abortus* data using the spatial binary probit model (1)-(4) by including an individual feature related to the individual species in *E*(*Y*_*sjl*_). For the three species elk, bison, and cattle, we included a common intercept *µ*_*e*_, *µ*_*b*_, and *µ*_*c*_ respectively. We also considered a simpler model that only includes one common intercept for all the species.

Although sample covariance is not the best estimator for true covariance when SNPs are dependent, we believed using all available SNPs data provides more insight into the rate of spread of *B. abortus* compared with using only an independent subset of the data. We estimated the parameters of different models using two sets of SNPs data: *set*_*full*_ and *set*_*tag*_. We compared results of the two different estimations using a 10-fold cross validation procedure as explained below.

To find which model under which estimation method works better in terms of out-of-sample prediction, we performed a 10-fold cross validation on the covariance matrices. We randomly divided the *N* individuals in *S* locations where genetic samples were collected into 10 equal-sized parts (folds). For each fold, we held out the observations related to one of these parts and estimated the parameters of each model from the remaining observations using the two data sets *set*_*full*_ and *set*_*tag*_. Then we used the estimated parameters to find Σ_***Y***_ (the model-based covariance matrix of the binary response) for each model.

Let 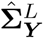 represent the estimated Σ_***Y***_ based on the landscape model, 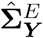 represents the estimated Σ_***Y***_ based on the exponential model, *ℋ*_*k*_ represent the individuals in the *k*^th^ hold-out part, and *N/ℋ*_*k*_ represent the set of all individuals excluding the hold-out part *ℋ*_*k*_. For the exponential model and each hold-out part *ℋ*_*k*_, let 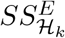 represents the sum of squared differences between the hold-out elements of 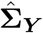 (the sample covariance matrix) and the hold-out elements of 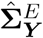. Let 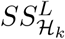 represents the same sum of squared differences for the landscape model. To make a fair comparison between the estimated parameters, 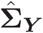 is calculated for the independent set of SNPs data in *set*_*tag*_. We calculate these values for each fold *k ∈ {*1, 2, …, 10*}* using the following formulas:

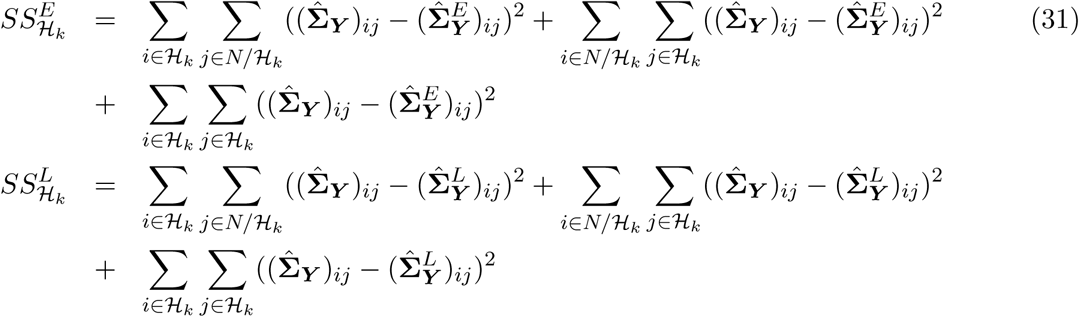

Figures (3a) and (3b) show more detail related to (31). In figure (3a), the yellow area (**A**) shows the elements of 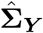 associated to locations in *N/ℋ*_*k*_ (non-hold-out elements), and the orange areas (**B**), (**C**), and (**D**) show the elements of 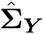 associated to locations in *ℋ*_*k*_ (hold-out elements). Figure (3b) shows the same thing for the estimated model-based covariance matrix 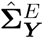 (or 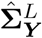). In the first double-sum of (31), we calculate the sum of squared differences between the elements of part (**B**) in figure (3a) and the elements of part (**B**) in figure (3b). The second double-sum calculates the sum of squared differences between the elements of part (**C**) in figure (3a) and the elements of part (**C**) in figure (3b). And in the third double-sum, we calculate the sum of squared differences between the elements of part (**D**) in figure (3a) and the elements of part (**D**) in figure (3b).

**Figure 3:**
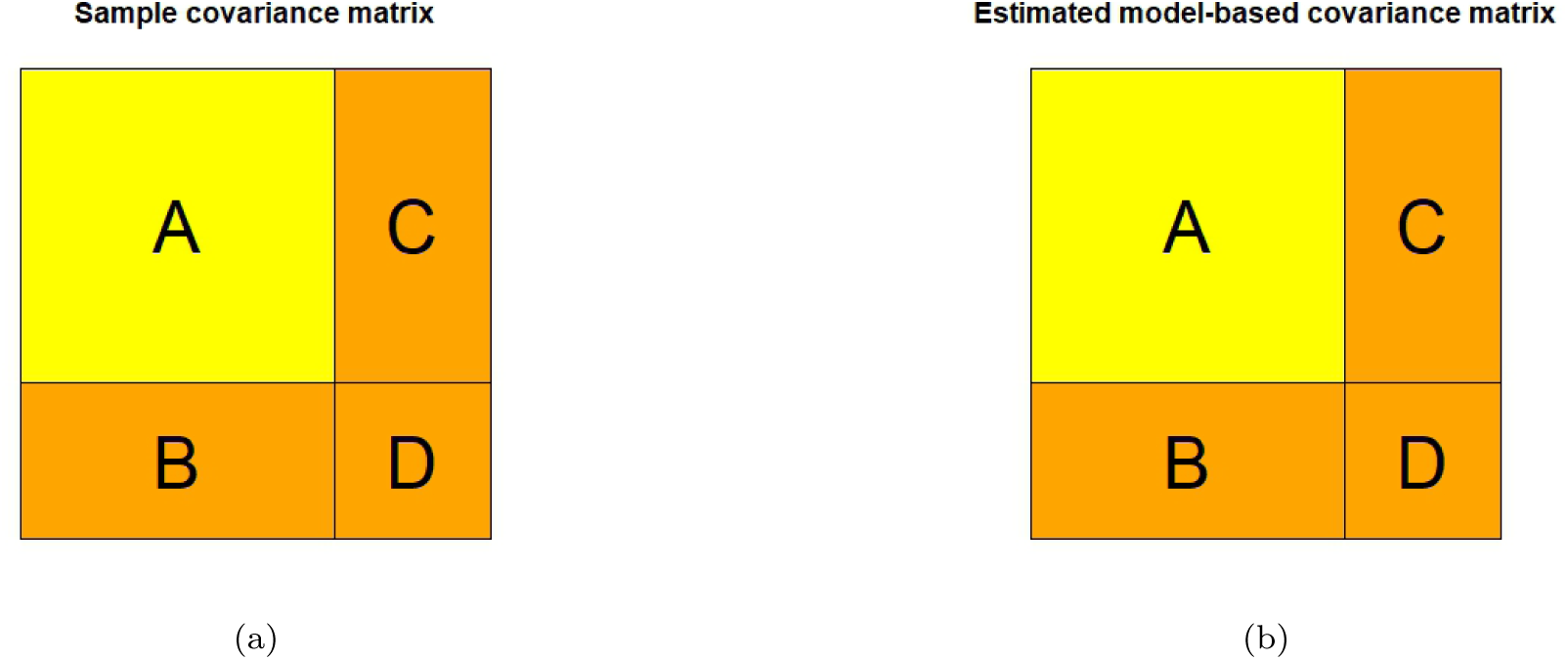
(a) Yellow area (**A**) shows the non-hold-out elements of 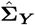. The orange areas (**B**), (**C**), and (**D**) show the hold-out elements of 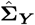 (b) Yellow area (**A**) shows the non-hold-out elements of 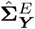 (or 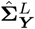). The orange areas (**B**), (**C**), and (**D**) show the hold-out elements of 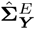 (or 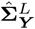).

Let *MSS*^*E*^ and *MSS*^*L*^ represent the mean of 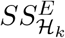 for *k* = 1, 2, …, 10 and the mean of 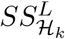 for *k* = 1, 2, …, 10 respectively. To compare different models under different methods of estimation, we compare *MSS*^*E*^ with *MSS*^*L*^ calculated as follows:

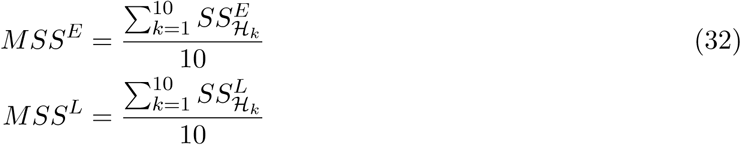

Table (3) represents *MSS*^*L*^ and *MSS*^*E*^ based on the estimated parameters for different models using two different sets of SNPs for both sample-based covariance approach (*COV*) and lorelogram-based approach (*LOL*).

**Table 3:** Summary of 10-fold cross validation error for landscape and exponential covariance models using all available SNPs (*set*_*full*_) and a subset of SNPs (*set*_*tag*_) chosen to be approximately independent of each other.

| Model |  | Set under which parameters are estimated |  |
| --- | --- | --- | --- |
| | | $set_{full}$ SNPs | $set_{tag}$ SNPs |
| COV | Exponential with common $\mu$ | 0.93389 | 0.01084 |
| | Exponential with $\mu_e, \mu_b, \mu_c$ | 0.88823 | 0.01101 |
| | Landscape with common $\mu$ and landscape covariates int, elev, and frst | 0.75521 | 0.01560 |
| | Landscape with $\mu_e, \mu_b, \mu_c$ and landscape covariates int, elev, and frst | 0.70522 | 0.01594 |
| | Landscape with common $\mu$ and landscape covariates int and elev | 0.76015 | 0.01545 |
| | Landscape with $\mu_e, \mu_b, \mu_c$ and landscape covariates int and elev | 0.70306 | 0.01579 |
| | Landscape with common $\mu$ and landscape covariates int and frst | 0.80770 | 0.01576 |
| | Landscape with $\mu_e, \mu_b, \mu_c$ and landscape covariates int and frst | 0.69771 | 0.01494 |
| LOL | Exponential with common $\mu$ | 42.06475 | 0.01660 |
| | Exponential with $\mu_e, \mu_b, \mu_c$ | 37.60402 | 0.01660 |
| | Landscape with common $\mu$ and landscape covariates int, elev, and frst | 135.13148 | 0.01660 |
| | Landscape with $\mu_e, \mu_b, \mu_c$ and landscape covariates int, elev, and frst | 145.67985 | 0.01660 |
| | Landscape with common $\mu$ and landscape covariates int and elev | 161.50350 | 0.01660 |
| | Landscape with $\mu_e, \mu_b, \mu_c$ and landscape covariates int and elev | 163.23229 | 0.01660 |
| | Landscape with common $\mu$ and landscape covariates int and frst | 148.18003 | 0.01660 |
| | Landscape with $\mu_e, \mu_b, \mu_c$ and landscape covariates int and frst | 147.83515 | 0.01660 |

For all the models, the results from the estimated parameters of tag SNPs set outperform the results from estimated parameters by using all available SNPs data in terms of out-of-sample prediction. This illustrates the importance of using independent loci to estimate the sample covariance matrix 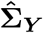. Appendix B shows that using dependent loci leads to a biased estimate of 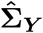. The estimation method *COV* outperforms *LOL* in all model settings. For the *COV* exponential model, including only one common intercept performs slightly better than the model with three different intercepts for three different species. For the *COV* landscape covariance, the model by including only three intercepts and two landscape covariates (intercept and percent of forested cells) provides better results compared to the other three landscape models. Between the two covariance models, the exponential model performs better than the landscape model by having a smaller value of *MSS*^*E*^ compared to the value of *MSS*^*L*^.

Based on the results from Table (3), we report the parameter estimates using the *COV* method for the exponential model with only one common intercept and the landscape model with three different intercepts and the two landscape covariates intercept and percent of forested cells from the data in the *set*_*tag*_ SNPs set. The point estimates for all parameters are obtained by numerically minimizing the squared error in (18) and (19). Our method of tag SNPs identified 386 clusters of SNPs, so we randomly picked one of the SNPs from each cluster as a representative of that cluster. However, both thinning the original data for construction of the tag SNPs set and random selection of clusters representatives caused removing many of *Y*_*sjl*_ = 1 from the data and made the constructed data very sparse. We calculated the 95% confidence intervals (CI) for the parameters of each model using an empirical bootstrap approach. For each model, we used 1000 bootstrap iterations and constructed the CIs from the estimated parameters of each bootstrap data set of size 386. All the computations are done on similar machines as the ones mentioned in section 4 and paralleled over 19 processors.

Table (4) represents the results of exponential covariance model:

**Table 4:**
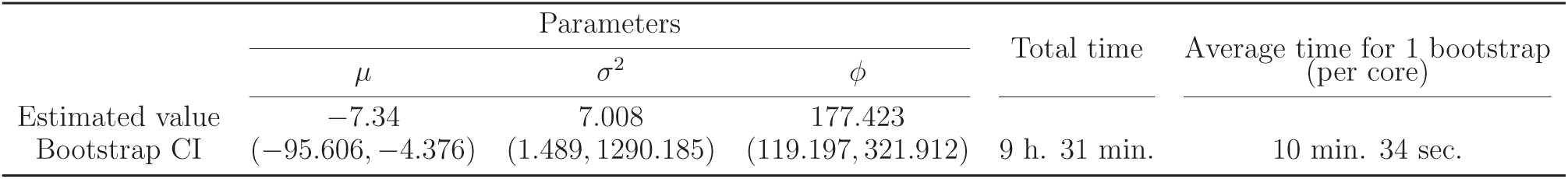
Estimated parameters and bootstrap CIs for *B. abortus* data using exponential covariance model and SNPs set *set*_*tag*_.

|  | Parameters |  |  | Total time | Average time for 1 bootstrap<br>(per core) |
| --- | --- | --- | --- | --- | --- |
| | $\mu$ | $\sigma^2$ | $\phi$ | | |
| Estimated value | -7.34 | 7.008 | 177.423 |  |  |
| Bootstrap CI | (-95.606, -4.376) | (1.489, 1290.185) | (119.197, 321.912) | 9 h. 31 min. | 10 min. 34 sec. |

Table (5) shows the results obtained by applying the landscape model to the *B. abortus* data. The best model included three different intercepts and the landscape covariates intercept with parameter *β*_0_, and percent of raster cells that are forested with parameter *β*_1_.

**Table 5:** Estimated parameters and bootstrap CIs for *B. abortus* data using landscape covariance model. *β*_0_ is the parameter associated with the intercept, *β*_1_ is the parameter associated with the percent of forested cells.

|  | Parameters |  |  |  |  | Total time<br>(per core) | Average time for 1 bootstrap |
| --- | --- | --- | --- | --- | --- | --- | --- |
| | $\mu_e$ | $\mu_b$ | $\mu_c$ | $\beta_0$ | $\beta_1$ | | |
| Estimated value | -4.034 | -3.965 | -4.229 | 3.136 | -2.026 |  |  |
| Bootstrap CI | (-4.138, -3.995) | (-4.234, -3.790) | (-4.396, -3.917) | (3.008, 3.323) | (-2.30, -1.82) | 15 h. 53 min. | 17 min. 47 sec. |

In both models, the value of *µ* is negative which implies that it is less likely to have a SNP than not to have a SNP in a specific location *s* and loci *l*. We note that our objective function depends on 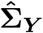, which is symmetric with respect to *µ*. Thus, the objective function is invariant to the sign of *µ*, and our approach can only estimate |*µ*|. Since SNPs are rare in the genetic data, we would expect to have a negative value of *µ*. Recovering a negative value for *µ*, which is more consistent with data, can be done by either constraining *µ* to be negative in the optimization algorithm, or by choosing the initial value of *µ* to be negative and small. Interpreting *ϕ* and σ^2^ is as straight forward for binary spatial models as it is for Gaussian spatial models. From the results, we see that as the distance increases to more than 200 kilometers, the correlation decays to 0.05 or less, which is also evident in figure (4b).

**Figure 4:**
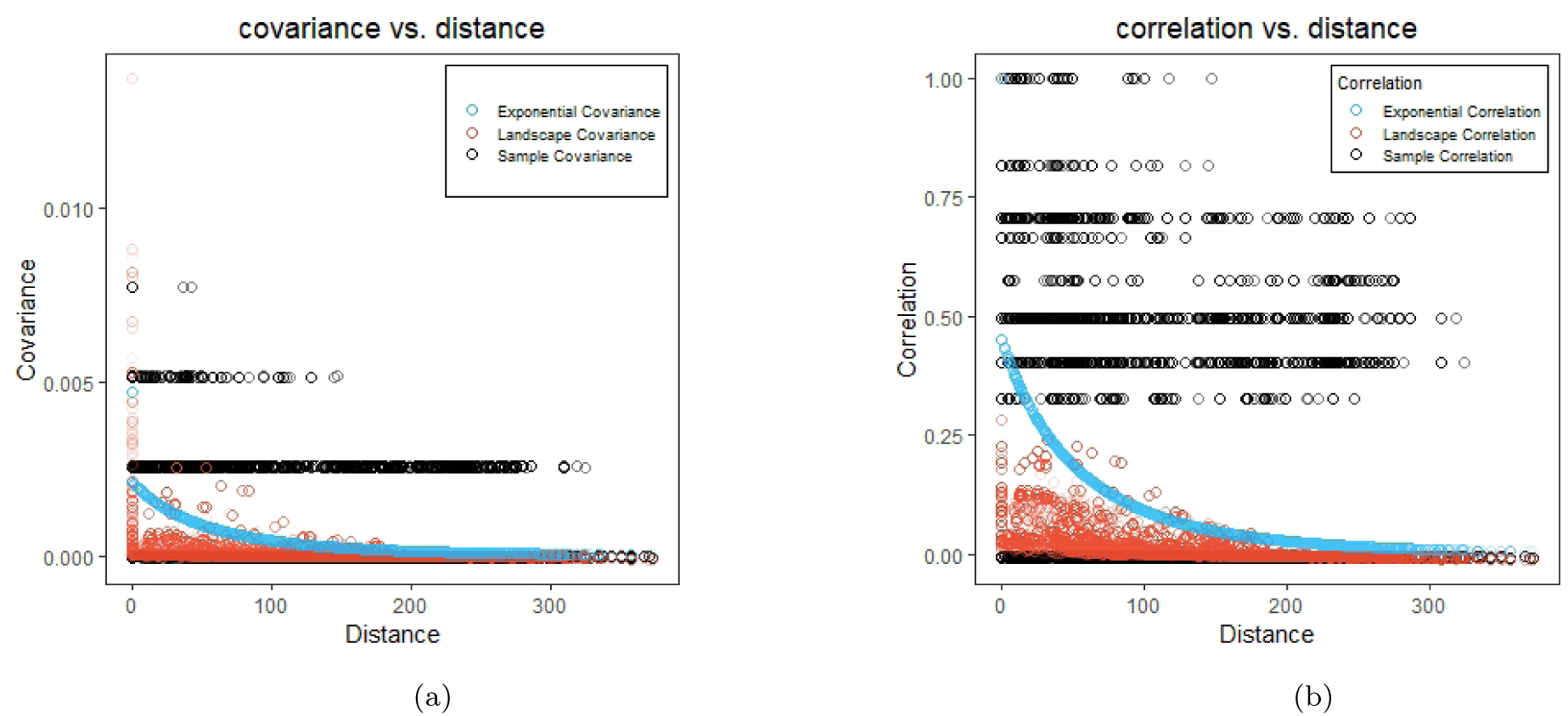
(a) The black dots represent the sample covariance, the red dots represent the covariance based on the landscape model, and the blue dots represent the covariance based on the exponential model. (b) The black dots represent the sample correlation, the red dots represent the correlation based on the landscape model, and the blue dots represent the correlation based on the exponential model.

For the landscape model, the negative value of *β*_1_ implies that the areas with lower percent of raster cells that are forested facilitates gene flow more than areas with higher percent. The average time needed to estimate one set of bootstrap parameters in the exponential model is around 10 minutes and 34 seconds, and is about 17 minutes and 47 seconds for the landscape model. Although performing 1000 bootstrap iterations sequentially would take several days, we can easily parallelize iterations on as many cores as possible and reduce the time significantly. To show how well each model describes the *B. abortus* data, we plotted the sample covariance of the tag SNPs data versus the distance between individuals and added the estimated Σ_***Y***_ in (16) for landscape and exponential models respectively. A similar plot for correlation versus distance was also generated. Figure (4) contains these two plots:

Figures (4a) and (4b) show that both models have some difficulties in capturing higher correlation between sites. Removing many of the SNPs data to construct a set of approximately independent tag SNPs set might be one of the reasons for these difficulties. The lack of fit in these two models might be because of missing model covariates or because we have considered the spatial locations of genetic samples to be fixed. Elk can move hundreds of kilometers over the course of a year, but the *B. abortus* data only include their last observed locations which may explain why locations with higher correlation in the data do not match with locations associated with higher correlation calculated by the models.

We check the validity of our cross validation approach by running two more simulations. The results, which are presented in the appendix C, show that our method selects the right model for independent simulated data. In addition, we perform a simulation study on both models by generating independent data set. In both studies, the true parameter values are estimated accurately and they are included in the bootstrap confidence intervals. The result of this simulation study can be found in the appendix D.

## 6 Discussion

In this paper, we investigated isolation by distance and isolation by resistance patterns in *B. abortus* data. We modeled the data using a GLMM with spatially referenced random effects. Due to the large size of the data, using common approaches in spatial statistics such as MCMC for inference is computationally taxing. We developed a novel method based on the properties of probit model and bivariate normal probabilities that computes the model covariance using numerical approximation. We based parameter estimation on minimizing the sum of the squared distance between the elements of the sample covariance and model-based covariance matrices. This sample covariance-based approach estimation for binary spatial models is the main contribution of this work. The form of the model covariance provided in (16) allows for computationally efficient evaluation of the model covariance as well as flexibility in using any covariance in the linear predictor of the spatial GLMM.

Our method can be applied to any kind of binary data which is modeled as in (1)-(4). This has been shown in the paper by analyzing two different covariance structures. We used an exponential covariance function in order to investigate the isolation by distance pattern in the *B. abortus* data. To investigate the isolation by resistance pattern, we considered a landscape covariance model. In this setting, our proposed method can be adapted to analyze spatially referenced genetic data under more general modeling framework. For example, we can specify different *µ*_*sj*_ for different locations or even model *µ*_*sj*_ based on some landscape features..

Our approach is based on the independence assumption of SNPs data. If the SNPs are dependent, the sample covariance is a biased estimator of the true covariance. When the independence assumption is violated, we demonstrated the possibility of choosing an independent subset of SNPs. However, by choosing independent tag SNPs, we may loose some important information in the data. Future work will include using a consistent covariance estimator when there are dependencies among loci. It is also possible that in some locations, the chance of seeing a SNP is higher than the other locations. Hence, specifying different *µ*_*sj*_ for different locations or modeling *µ*_*sj*_ by including land-scape features might improve our model in explaining the patterns in the *B. abortus* data. In the case of *B. abortus* data, it might be more appropriate to consider a spatio-temporal covariance matrix. The data is collected from 1985 to 2013, so considering the time in the model might improve our analysis.

To improve the efficiency of the current estimation procedure, future work will consider using weighted least squares instead of our current approach. Fitting our model by a weighted least squares approach would be similar to fitting variogram models using this approach (Cressie, 1985). In our case, weights could come from the large-sample variance of the elements of the vectorized sample covariance matrix (Bilodeau and Brenner, 1999). An alternative approach to improve computing would be to consider parallel MCMC approaches (e.g. Bardenet et al. (2017)), which would take advantage of the independence of SNPs.

In this work, we used the latent variable method of Albert and Chib (1993) to represent the binary probit model. Polson et al. (2013) proposed an exact and simple method for fully Bayesian inference in logit model which appeals to Poly-Gamma distributions. In Future, we will extend our work to binary logit model and compare the result of Bayesian analysis of logit model with our current results.

Our simulation study compares the performance of our method with MCMC which is most commonly used for spatial binary data as well as lorelogram-based approach which is another approach for analyzing spatial binary data. However, we found the lorelogram numerically unstable and future work could explore more accurate and stable numerical approaches for lorelogram computations. There are some papers in the literature which suggested using Generalized estimating equation (GEE) (Albert and McShane, 1995; Augustin et al., 2005; Carl and Kuhn, 2007; Dormann et al., 2007). Future work will extend our comparison to the less common GEE methods.

As spatial data collection becomes easier, large spatial data sets will be more available to researchers. Our approach illustrates the computational efficiency of estimation approaches based on sample covariance of binary spatial data.

## 7 Acknowledgments

Funding: This work was partially supported by the United States Geological Survey [grant number G16AC00055].

### Appendix A Details of MCMC algorithm for simulation study

In section 3.1, we introduced the Bayesian approach for estimating model parameters. Algorithms (1) and (2) provide additional detail on the MCMC algorithm for the exponential model.

#### Algorithm 1 MCMC algorithm for SNP data

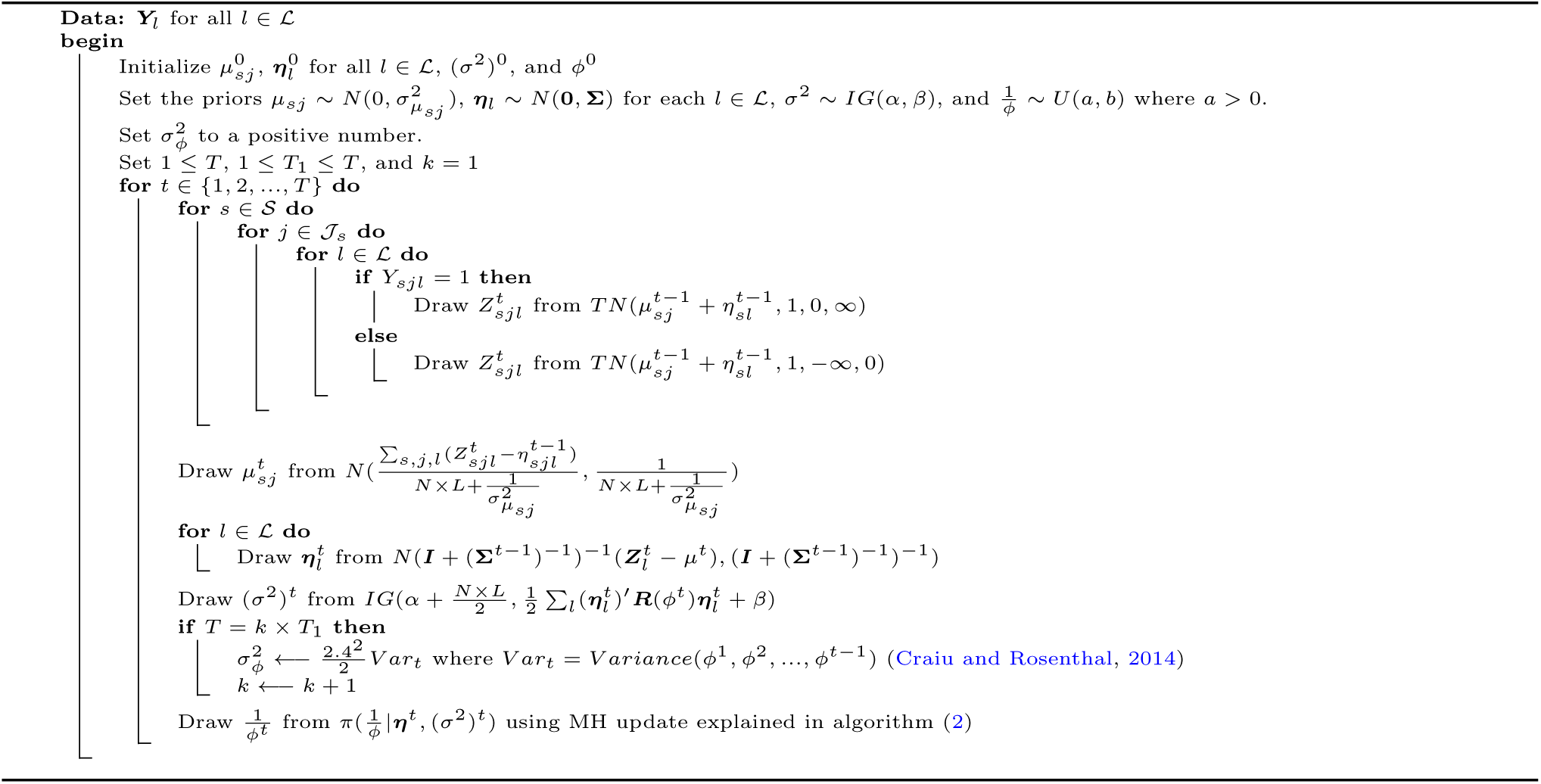

### Appendix B Bias of the sample covariance as an estimator of true covariance

Let ***µ*** be a vector of size *N* with elements *µ*_*sj*_. Following our model (1)-(4), for any *l ∈ ℒ*:

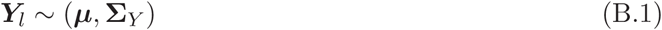

#### Algorithm 2 MH update for parameter ϕ

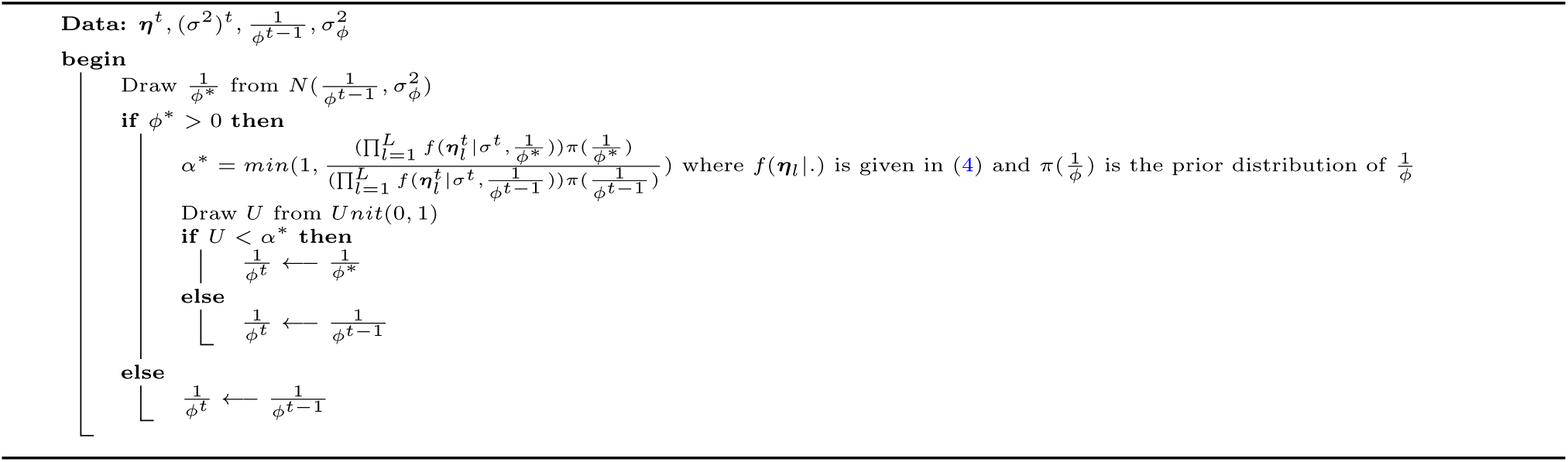

If ***Y***_*l*_ and ***Y***_*l′*_ for the two loci *l* and *l′* be dependent, then:

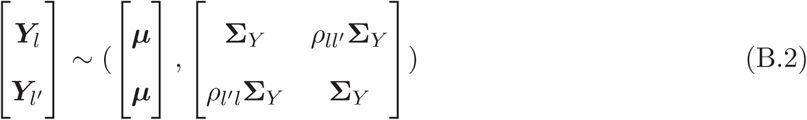

where *ρ*_*ll′*_ = *ρ*_*l′l*_ is the correlation between SNPs in loci *l* and *l′*.

Let 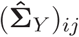 represents the sample covariance of ***Y*** for *i ∈ 𝒥*_*s*_ and *j ∈ 𝒥*_*s′*_which can be calculated as follows:

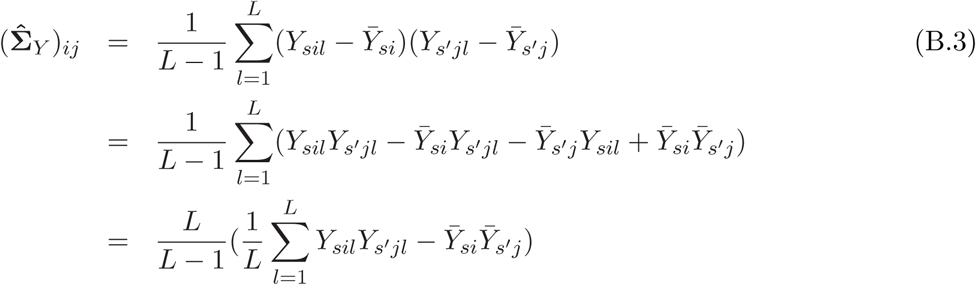

To find the bias, we need to calculate 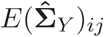 as follows:

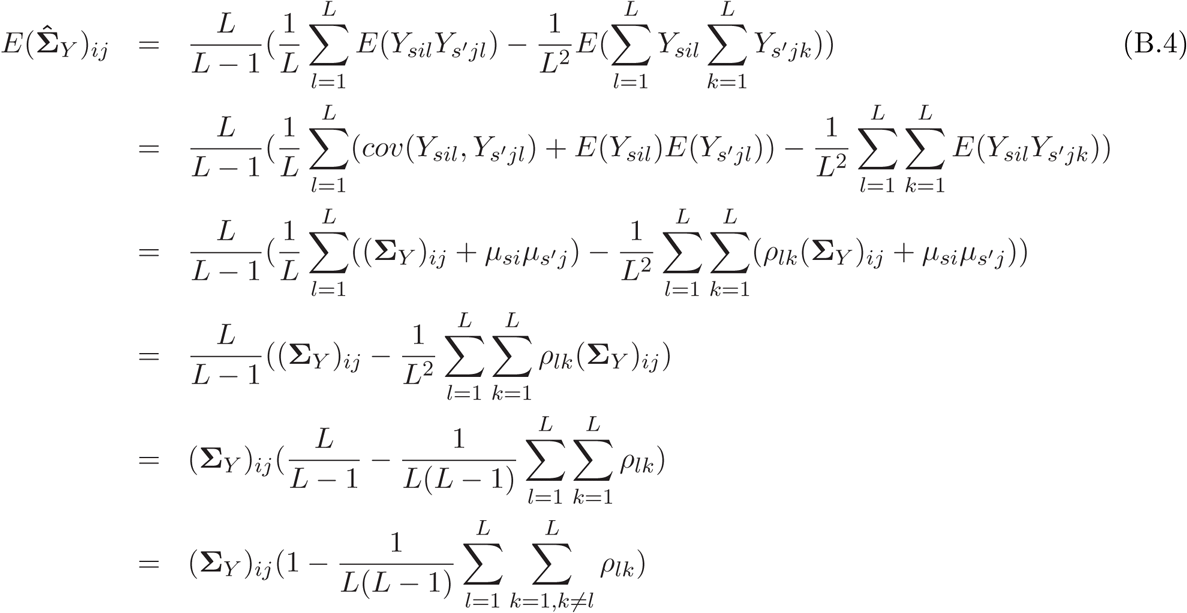

Therefore, sample covariance is a biased estimator when the SNPs are not independent.

### Appendix C Cross validation for simulated data using both covariance models

To check the validity of our cross validation approach, we ran two more simulations. In the first simulation, we generated data according to the exponential covariance model. To generate the data, we used the locations in the *B. abortus* data and calculated the distance matrix *D*. We set the values of *µ*_*sj*_, σ^2^, and *ϕ* to some specific values and simulated ***Y***_*l*_ for all *l ∈ ℒ* according to the exponential covariance model. Then we used the same cross validation approach explained in Section 5 on the simulated data to compare the two models. Note that for the landscape covariance method, we used the intercept and elevation rasters in the *B. abortus* data.

Table (C.1) represents the results for the exponential simulated data. The true parameter values are *µ*_*sj*_ = *µ* = −5, σ^2^ = 4, and *ϕ* = 50.

**Table C.1:**
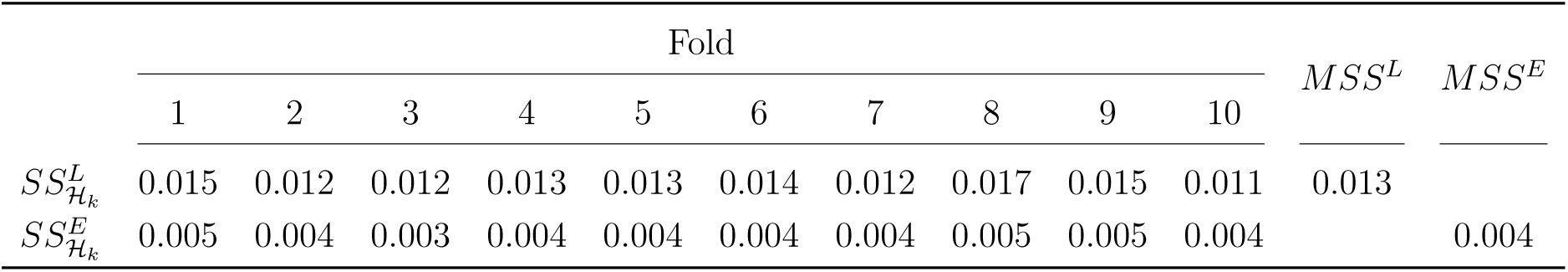
Summary of 10-fold cross validation results for landscape and exponential covariance models

Since we simulated the data using the exponential model, we would expect our cross validation approach selects the exponential model. From table (C.1) results, we confirmed that our model selection approach picks the right model.

In the second simulation, we generated the data according to the landscape covariance model. We used the same locations as *B. abortus* data. To simulate the data, we used the intercept and elevation rasters from the *B. abortus* data and set the model parameters values to some specific numbers. Then we generated ***Y***_*l*_ for all *l ∈ ℒ* according to the landscape covariance model.

Table (C.2) represents the results when *µ*_*sj*_ = *µ* = −3.9687, *β*_0_ = 1.5825, and *β*_1_ = 0.2874.

**Table C.2:** Summary of 10-fold cross validation results for landscape and exponential covariance models

| | Fold | | | | | | | | | | $MSS^L$ | $MSS^E$ |
| --- | --- | --- | --- | --- | --- | --- | --- | --- | --- | --- | --- | --- |
|  | 1 | 2 | 3 | 4 | 5 | 6 | 7 | 8 | 9 | 10 |  |  |
| $SS_{\mathcal{H}_k}^L$ | 0.017 | 0.021 | 0.021 | 0.019 | 0.017 | 0.021 | 0.024 | 0.018 | 0.017 | 0.023 | 0.020 | |
| $SS_{\mathcal{H}_k}^E$ | 0.219 | 0.418 | 0.397 | 0.328 | 0.376 | 0.444 | 0.441 | 0.247 | 0.334 | 0.411 | | 0.362 |

Similar to the previous results, our cross validation method selects the right model which is the landscape covariance model.

### Appendix D Simulation study on both exponential and landscape models

To show our method is capable of estimating the true parameter values in different settings, we perform a simulation study using both exponential and landscape models. We generate data with the same size of the *B. abortus* data which includes SNPs in *L* = 1463 loci from *N* = 237 individuals. We find the parametric bootstrap confidence intervals in the same way as explained in Section 4.

Tables show the results for exponential model and landscape model respectively.

Both tables show that our method is able to estimate the model parameters accurately and the bootstrap confidence intervals include the true parameters values.

**Table D.1:**
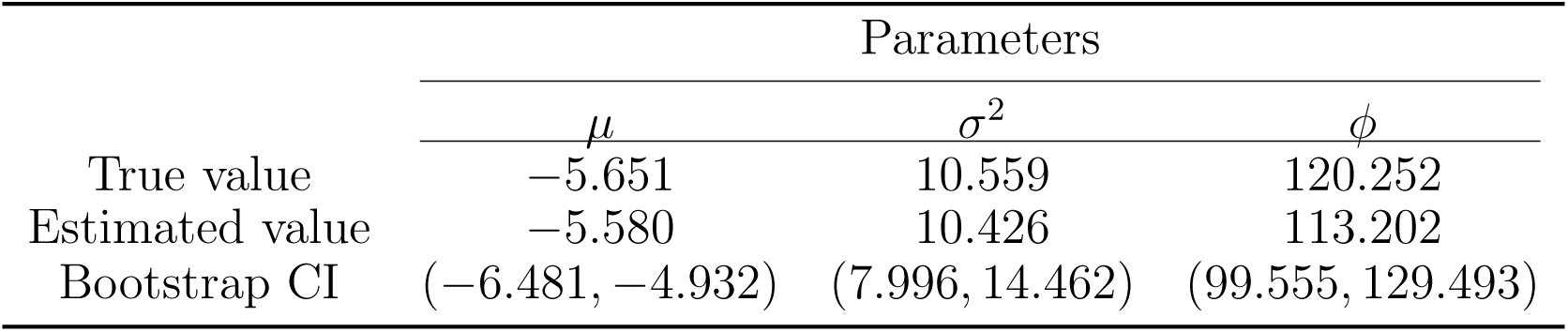
Simulation results when data is generated based on the exponential covariance model.

|  | Parameters |  |  |
| --- | --- | --- | --- |
| | $\mu$ | $\sigma^2$ | $\phi$ |
| True value | -5.651 | 10.559 | 120.252 |
| Estimated value | -5.580 | 10.426 | 113.202 |
| Bootstrap CI | $(-6.481, -4.932)$ | $(7.996, 14.462)$ | $(99.555, 129.493)$ |

**Table D.2:** Simulation results when data is generated based on the landscape covariance model.

|  | Parameters |  |  |  |
| --- | --- | --- | --- | --- |
| | $\mu$ | $\beta_0$ | $\beta_1$ | $\beta_2$ |
| True value | -3.565 | 1.684 | 0.255 | 0.062 |
| Estimated value | -3.454 | 1.693 | 0.222 | 0.179 |
| Bootstrap CI | $(-3.671, -3.285)$ | $(1.506, 1.944)$ | $(0.063, 0.354)$ | $(-0.398, 0.621)$ |

